# A Robust and Generalizable Low-Input Spatial Proteomics Workflow Enabling Deep Proteome Coverage

**DOI:** 10.64898/2026.07.16.738934

**Authors:** T. Courtellemont, R. Hamelin, F. Armand, R. Dornier, J. Sordet-Dessimoz, MP. Pavlou

## Abstract

Spatial proteomics has rapidly emerged as a powerful approach for deciphering the molecular architecture of complex tissues, offering critical insights into how local proteome composition shapes tissue function. In this work, we aimed to broaden access to mass spectrometry-based spatial proteomics by putting in place a pipeline relying exclusively on instrumentation commonly available in standard histology, imaging, and proteomics facilities. The resulting workflow integrates high-resolution whole-slide imaging, image-based region selection, streamlined low-input sample processing, and a high-sensitivity method optimized on the Orbitrap Exploris platform. To demonstrate the potential of this approach, we applied it to the choroid plexus (ChP), a highly specialized but understudied brain structure whose spatial molecular organization remains poorly characterized. Analysis of ChP samples from three brain compartments across mice of different age and sex quantified more than 8,000 protein groups across the dataset from minimal input material, providing unprecedented proteome-wide resolution of ChP spatial heterogeneity. Our findings establish a foundational spatial proteomic atlas of the ChP and highlight new opportunities for investigating its roles in aging, neurological disease, and central nervous system infection.

**Graphical abstract:** 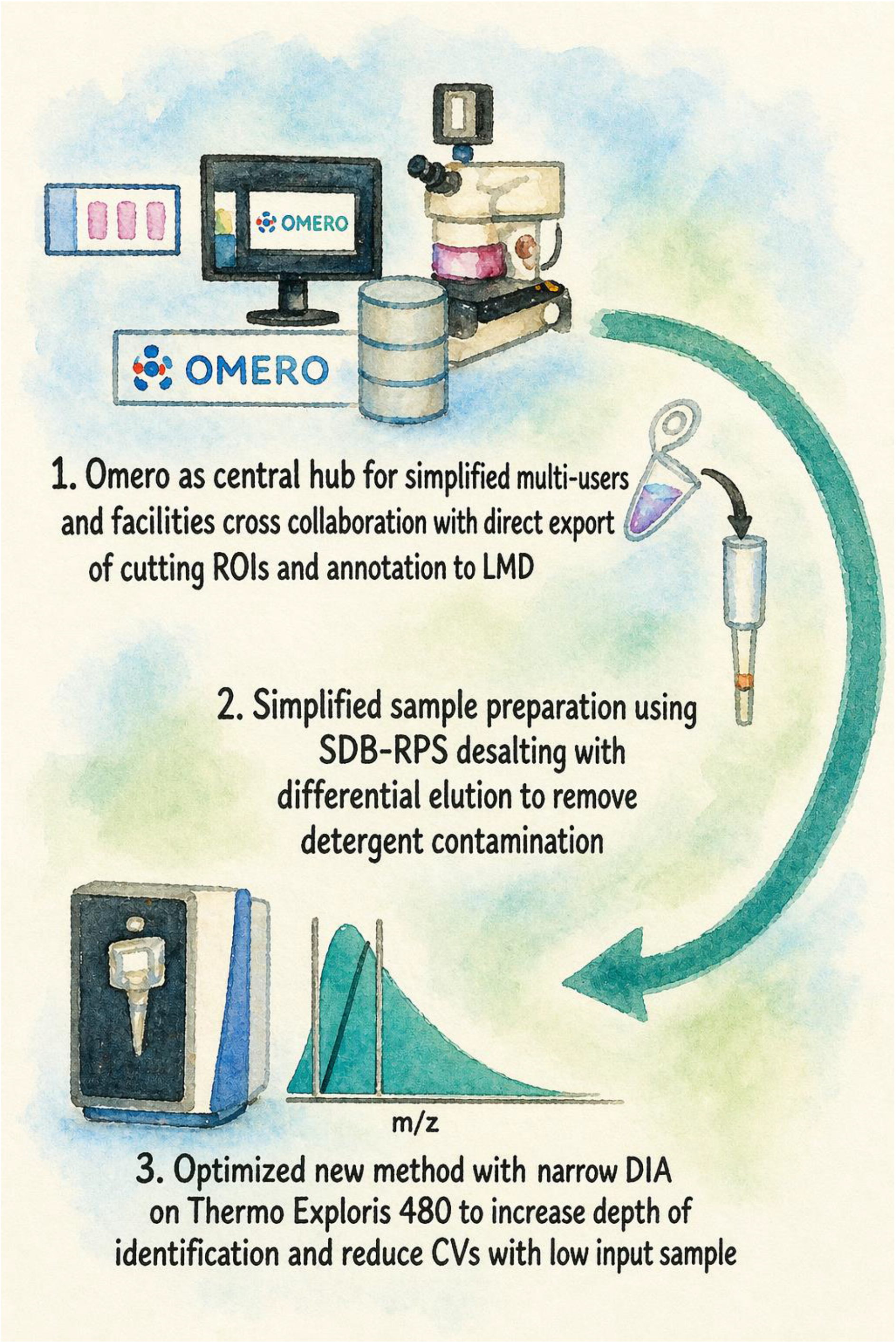

## Introduction

Core facilities serve as ambassadors for their technological focus. They provide sophisticated instrumentation and deep expertise on technologies that individual labs cannot readily adopt due to cost and the need for specialization. For example, expert-level proficiency in mass spectrometry-based proteomics typically requires 5-10 years of focused training and hands-on experience, reflecting the multidisciplinary nature of the field. Platforms must serve a broad range of research questions in a time and cost efficient manner. They should also scout for new workflows and, when relevant, adopt and offer them. The field is pushed forward typically by leading labs which usually have access to the latest and most advanced instrumentation. However, instrument acquisition and implementation carry a significant investment and cannot be routinely performed by academic institutions. In such cases, the performance of existing instruments must be fine-tuned to provide the best possible results.

The importance of tissue architecture and heterogeneity has been recognized for a long time, but it has been rediscovered over the last five years at the omics level. These advancements enable the study of an impressive number of analytes at both the mRNA and protein levels with the community clearly acknowledging the significance of these novel methodologies^1,2^

Spatial proteomics (SP) includes a wide range of technologies that differ in scale, resolution, and sensitivity. Current spatial strategies fall broadly into two categories: antibody-based using microscopy or mass spectrometry as readout and target agnostic exploratory mass spectrometry-based approaches. High-dimensional antibody-based imaging platforms such as Lunaphore COMET and Akoya PhenoCycler enable simultaneous spatial profiling of dozens of proteins at single-cell resolution, making them well suited for targeted marker screening^3,4^. MALDI-IHC^5^ serves as a powerful hybrid approach combining the spatial and morphological advantages of traditional immunohistochemistry with the comprehensive analytical capabilities of mass spectrometry-based detection and it offers another powerful workflow for studies where the protein targets are known. However, for exploratory studies aiming to generate new hypotheses, unbiased approaches are more favorable. Tissue microdissection guided by hematoxylin-eosin (H&E), immunohistochemical (IHC) or immunofluorescent (IF) staining, coupled with ultrasensitive mass spectrometry, has massively accelerated the profiling of thousands of proteins while preserving spatial information. Over the past six years, several workflows have been published that combine optimal sample handling to minimize losses with advanced mass spectrometry methods for sensitive, reproducible, and high-throughput analysis including Deep Visual Proteomics (DVP)^6^, FAXP^7^ and nanoPOTS^8^.

Recognizing the significant value and potential impact of these workflows for advancing research across life science community, the proteomics core facility (PCF) of the School of Life Sciences at EPFL partnered with the histology (PHT) and imaging (BIOP) platforms to successfully establish and implement a comprehensive spatial proteomics workflow which takes advantage of and builds upon existing instrumentation already available within the School of Life Sciences. The developed workflow integrates histological sample preparation, high-resolution bioimaging, and advanced data-independent acquisition (DIA) mass spectrometry. To showcase the potential of this method, we performed spatial proteomic profiling of the choroid plexus; a highly compartmentalized, specialized yet understudied brain structure; from mice of different age and sex.

## Results

To establish a robust and reproducible spatial proteomics workflow, we developed an integrated pipeline that leverages standardized histology, semi-manual laser microdissection and liquid chromatography mass spectrometry (Figure 1). The workflow is designed to be hardware-agnostic, utilizing common laboratory infrastructure to ensure broad utility across different research environments. Our pipeline accommodates a diverse range of starting materials, including animal models, human clinical biopsies, and 3D-cultured organoids. Below, we present the developments in the different steps of the workflow. In box 1, we provide detailed information on parameters which are typically emitted from papers but remain critical for reproducing the workflow.

**Figure 1.**
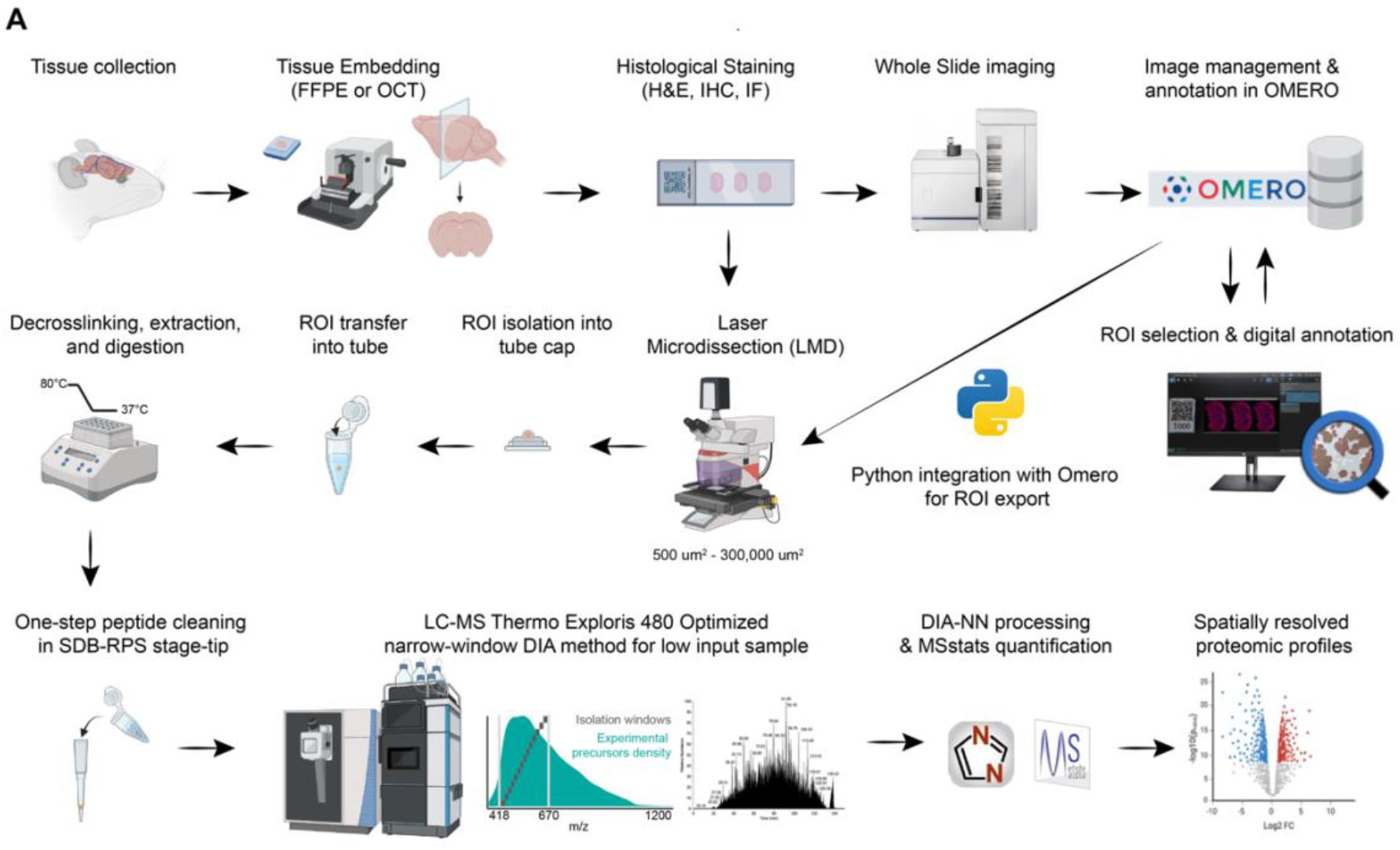
Integrated spatial proteomics workflow from tissue collection to bioinformatic analysis. Schematic overview of the spatial proteomics pipeline.

### Tissue microdissection

Specimens can be preserved either in formalin-fixed paraffin-embedding (FFPE) or optimal cutting temperature (OCT) compound for fixed and fresh-frozen applications, respectively. To ensure any potential hits could be independently verified, we implemented a parallel sectioning protocol. Serial sections are collected alternately onto glass microscopy slides for high-resolution imaging and PEN-membrane slides for Laser Capture Microdissection (LCM). This strategy allows precise correlation of proteomic signatures with specific histological features identified in adjacent sections. Following sectioning, slides undergo histological staining; primarily Hematoxylin and Eosin (H&E), though the workflow is compatible with Immunohistochemistry (IHC) and Immunofluorescence (IF) for marker-guided isolation. Digitized whole-slide images are managed via the OMERO platform^9^, an open-source environment for bio-image data management. A key feature of our pipeline is the integration of a custom Python script that bridges the gap between digital annotation and physical microdissection that could be linked directly to OMERO. User-defined regions of interest (ROIs), along with three border calibration points, are annotated in QuPath^10^, an open-source program that is well established for visualising and analysing Whole Slide Imaging (WSI) data, and are subsequently stored in OMERO. Their spatial coordinates are then directly exported to the LCM system, thereby reducing manual intervention and minimizing subjectivity in ROI selection. Additionally, OMERO can be coupled with external imaging software such as FIJI^11^ or QuPath for image enhancement tasks, including segmentation and cell counting. Tissue areas are isolated via LMD and captured directly onto tube caps.

### Sample preparation

Samples are transferred from the cap to the bottom of the tube using acetonitrile, then dried in a SpeedVac before sample preparation. Inspired from digestion protocols used for nearly single or single cell proteomics^12,13^, we aimed for a single tube reaction using the mass spectrometry compatible surfactant *n*-dodecyl-β-D-maltoside (DDM). For FFPE samples, efficient processing required a heat-induced decrosslinking step combined with detergent-assisted extraction. Tissue dissections were first incubated at 80 °C for 90 min in 300 mM Tris-HCl, pH 8.0, containing 0.2% DDM, followed by progressive cooling to 37 °C. Digestion was then initiated by adding an equal volume of a concentrated digestion mixture. This enabled decrosslinking, protein solubilization, reduction/alkylation, and enzymatic digestion within a single low-volume workflow (Figure 1).

DDM is a well-documented detergent that solubilizes membrane proteins enabling efficient cell lysis and does not interfere with peptide identification as it elutes at high organic solvent concentration. However, DDM is still introduced into the mass spectrometer, which may lead to progressive contamination and interference with liquid chromatography over time. Moreover, our micro-ROI strategy requires relatively higher detergent concentrations compared to single-cell approaches. To minimize DDM introduction in the system, we evaluated the capacity of three desalting formats namely homemade stage-tips^14^ with C18 or SDB-RPS (Styrene Divinyl Benzene-Reverse Phase Sulfonate) phase-transfer membranes and commercially available Evotips^15^ to remove DDM while maintaining reproducible peptide recovery. To assess the reproducibility of peptide recovery among the different devices under baseline conditions, we first used commercial HeLa peptide mix (10ng) without DDM (Figure S1A-C). All three devices provided comparable results in terms of peptide recovery and protein group identification with the homemade stage-tips yielding slightly higher proteome coverage compared to commercial tips. In terms of reproducibility, homemade stage-tips consistently outperformed commercial tips with median Coefficient of Variation (CV) values of 3.9% and 4.3% for C18 and SDB-RPS respectively compared to 7.3% for Evotips (Figure S1C). We then produced HeLa peptide mix samples supplemented with 0.02% DDM, performed the desalting with the three different devices and analysed the samples via liquid chromatography mass spectrometry (LC-MS). We observed a cleaner baseline and a more consistent peptide elution profile in the SDB-RPS treated samples particularly in the higher organic elution part of the chromatographic gradient where DDM contaminants typically co-elute. The mass spectra confirmed that SDB-RPS effectively mitigates the presence of high-abundance contaminant clusters (e.g. m/z = 528.33; 1021.61) which are more prominent in samples desalted with C18 phase (Figure S1D-E). Therefore, desalting via SDB-RPS stage-tips was selected for the pipeline as it provides the most robust solution for transitioning from detergent-heavy lysis to high-sensitivity MS analysis without sacrificing proteomic depth or data reproducibility.

### LC-MS method development

A major focus of this work was to establish a data independent acquisition (DIA) strategy for low-input samples that could be implemented on conventional mass spectrometry platforms. Such a method requires a fine balance between sample throughput and optimal depth, accuracy, and reproducibility of protein quantification. With this in mind, we sought to push the performance of the Exploris 480, a widely used instrument whose acquisition speed remains more limited than that of newer-generation mass spectrometers. In the context of this study, which aims to analyse several dozen proteomic samples with a wide dynamic range, a 120-minute active gradient length used in numerous published studies was selected as an appropriate compromise^16–18^. To define the full set of peptides and protein groups that can be experimentally identified under our conditions using such a gradient and instrument, we employed a widely used acquisition strategy^19^ that scans a broad precursor m/z range, providing an unbiased view of the experimental distribution of identifiable precursors when analysed with DIA-NN. This acquisition was initially performed using a relatively large amount of material (300 ng HeLa digest) to facilitate robust detection of precursors that are expected to be identifiable, prior to a final injection of amounts resembling spatial proteomics samples.

This initial step allowed us to identify approximately 170,000 precursors across the four technical replicates. The distribution of identified precursors across the m/z range is shown in Figure 2A, where, as expected, we observed an uneven distribution throughout the mass range. Currently, most MS methods concentrate the data acquisition within a shorter m/z window^20,21^. Reducing the initial scan range by a factor of two (420-980) still allows coverage of more than 90% of the precursors identifiable under our experimental conditions. Numerous studies have demonstrated the benefit of focusing acquisition on m/z regions that contain high density peptide population and our approach follows this rationale^21,22^. Based on the precursor m/z ion density observed in our data, we defined several acquisition windows spanning 560, 392, 252, and 196 m/z, which respectively encompass 90%, 76%, 55%, and 45% of the total precursors identified in the initial broad m/z survey (Figure 2A).

**Figure 2.**
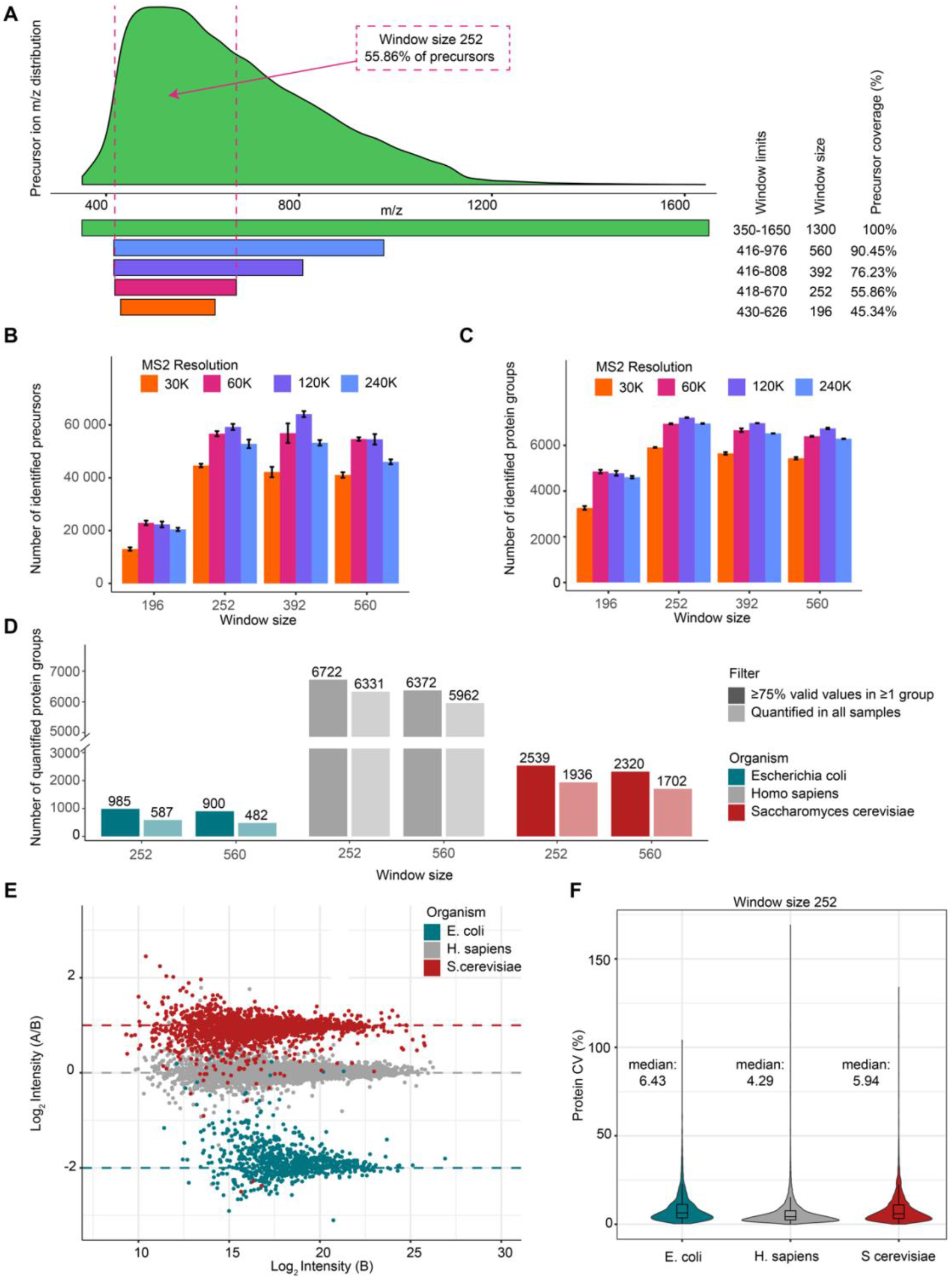
Optimization of DIA window size and MS2 resolution for best proteome coverage and quantitative performance. **(A)** Distribution of precursor ion *m/z* values across the analyzed dataset. Horizontal bars indicate the precursor *m/z* ranges evaluated for different DIA window size and the corresponding percentage of precursors covered by each range. **(B, C)** Effect of DIA window size and MS2 resolution on proteome coverage using 10 ng HeLa digest. Bar plots show the number of identified peptides **(B) and identified** protein groups **(C)** across different precursor *m/z* ranges and MS2 resolutions. Error bars indicate the standard deviation across technical replicates. **(D)** Quantitative depth obtained with the 252 and 560 *m/z* acquisition ranges in a three-proteome benchmark sample. Bars show the number of quantified protein groups for *E. coli*, *H. sapiens*, and *S. cerevisiae*. Dark bars represent protein groups with at least 75% valid values in at least one organism group, whereas light bars indicate protein groups quantified in all samples. Numbers above bars indicate the corresponding protein-group counts. **(E)** Quantitative accuracy of the selected 252 *m/z* acquisition method in the three-proteome benchmark. The scatter plot shows measured log₂ fold change between mixtures A and B as a function of mean log₂ intensity in mixture B for each quantified protein group. Dashed lines indicate the expected organism-specific log2 fold changes. **(F)** Quantitative reproducibility of the selected 252 *m/z* acquisition method. Violin plots with overlaid boxplots show the distribution of protein-level coefficients of variation (CVs) for each organism. Median CV values are indicated above each violin. Only protein groups with at least 75% valid values in at least one organism group were included in panels D–F.

Given the limited peptide input from spatial proteomics samples, we systematically optimized acquisition parameters to maximize sensitivity by varying precursor isolation window widths, injection times, and MS resolution settings.

To assess this across the four previously defined precursor m/z windows, we conducted a series of experiments in which a 10 ng HeLa digest was injected. MS spectra were acquired using DIA methods with varying precursor isolation window widths and MS resolutions, combined with different ion injection times, while maintaining constant scan cycle time (Supp Table 1) thereby preserving robust sampling while ensuring consistent and comparable quantitative performances. The progressive increase in resolution and accumulation times improves peptide and protein group identification across the three widest precursor windows (Figure 2B). An optimum for peptide and protein identification was observed for the method at 120k resolution. The highest number of identified peptides (64,109) was obtained with a 392 Da window, whereas the greatest analytical depth in terms of protein identifications (7,213) was achieved with a 252 Da window (Figure 2B-C).

With the objective of maximizing the number of identified protein groups, the 120k method with a 252 Da precursor window was selected for our pipeline. Pairwise correlation and coefficients of variation are widely used metrics for evaluating quantification performance. However, these metrics are primarily capturing precision rather than accuracy. To assess the suitability of our best-performing method for spatial proteomics, we evaluated proteome depth alongside quantitative accuracy and precision using a 10 ng three-proteome benchmark sample. Across the methods tested in our current study, 120k resolution methods generally outperformed the other acquisition strategies (Figure 2B-C). We then compared our 252 Da window acquisition with a more conventional MS acquisition method using a 560 Da window both using a similar 120k resolution. Similarly, to the 10 ng HeLa benchmark, our optimized method increases proteome depth across the three organisms (Figure 2D) and improved quantification reproducibility across four technical replicates while maintaining low protein quantification coefficients of variation (below 7% for spiked *E. coli* and yeast, and below 5% for the human background proteome) (Figure 2E-F & Figure S2B). Protein quantification accuracy for both methods was assessed as the percentage error between measured and theoretical fold changes^23^. The distributions of percentage errors were similar for E. coli proteins across the 2 methods but for human and yeast proteins exhibited slightly improved performance, with consistently lower error rates. (Figure S2A-C)

Together, these findings demonstrate that we optimized a sensitive microproteomic DIA method (SM-DIA) for low-input proteomics by refining precursor m/z windows sizes and acquisition parameters. This approach maximizes proteome depth while maintaining quantitative accuracy and precision, enabling robust and highly sensitive analyses.

### Application of optimized pipeline on LCM tissues of varying size and tissue types

To validate the performance and sensitivity of our pipeline on biological specimens, we applied the workflow to FFPE liver tissue sections (4 µm thickness) (Figure S3A). We performed serial dissections of circular ROIs with surface areas of 1,000 µm^2^, 3,000 µm^2^, 6,000 µm^2^, 12,000 µm^2^, 25,000 µm^2^, 50,000 µm^2^, and 100,000 µm^2^, which roughly correspond to a cell volume equivalent of 2, 6, 8, 14, 26, 45, and 84 measured hepatocytes, respectively (Figure S3C). The resulting data demonstrated a high degree of sensitivity, robustness and scalability across the entire titration range (Figure S3B). At the lowest input level of 1,000 µm^2^, representing closely a single hepatocyte, we successfully quantified over 1,500 protein groups and 7,500 precursors. As the dissected area increased, we observed a steady, near-linear increase in proteomic depth, reaching approximately 5,500 quantified protein groups and 40,000 quantified precursors at the 100,000 µm^2^ scale. The experiment was also replicated using fresh-frozen OCT-embedded tissue sections (10 µm thickness) microdissected with a PALM Zeiss microscope (Figure S3E). After accounting for differences in tissue thickness (10 µm vs 4 µm), the results were highly consistent with those obtained from the original samples (Figure S3F). Importantly, comparable coefficients of variation (CVs) were observed between both approaches, suggesting that the FFPE decrosslinking workflow does not substantially impair protein group identification or quantification reproducibility (Figure S3D-G).

To ensure that our pipeline is applicable to different specimens we performed spatial analysis of four different organs, namely the liver, spleen, brain and intestine (Figure 3A). For each FFPE tissue slice we dissected 4 sections of 100 000µm^2^ and followed the optimized pipeline (in terms of preparation and LC-MS analysis). We were able to consistently identify over 6,000 protein groups for each tissue (Figure 3A) and low protein coefficient of variation (ranging from 9.2% to 17.7% per group), also we observed a strong segregation of the sections coming from the same tissue (Figure 3B-C). Although most quantified protein groups were common across tissues, we quantified numerous differentially abundant protein groups with strong tissue specificity (Figure 3D). For the liver and spleen, the vast majority of these tissue-enriched protein groups were also annotated as tissue-specific or tissue-enriched in the Human Protein Atlas, providing additional validation of the data quality (Figure 3E). For the brain and intestine, we quantified a similar number of tissue-enriched protein groups; however, a substantial fraction was not classified as tissue-specific in the Human Protein Atlas. This likely reflects the more restricted set of tissues examined in our study, which enables the detection of proteins that are selectively enriched within this tissue panel but not necessarily tissue-specific across the broader range of tissues represented in the Human Protein Atlas. To further assess the biological relevance of the tissue-enriched proteins that were not annotated in the Human Protein Atlas, we performed tissue-term enrichment analysis using STRING-DB^24^. This analysis revealed significant enrichment for tissue-relevant terms in both the brain and intestine but also for the liver datasets (N.A. for Spleen), indicating that these proteomics-derived candidates are consistent with the expected biology of the corresponding tissues (Figure 3F).

**Figure 3.**
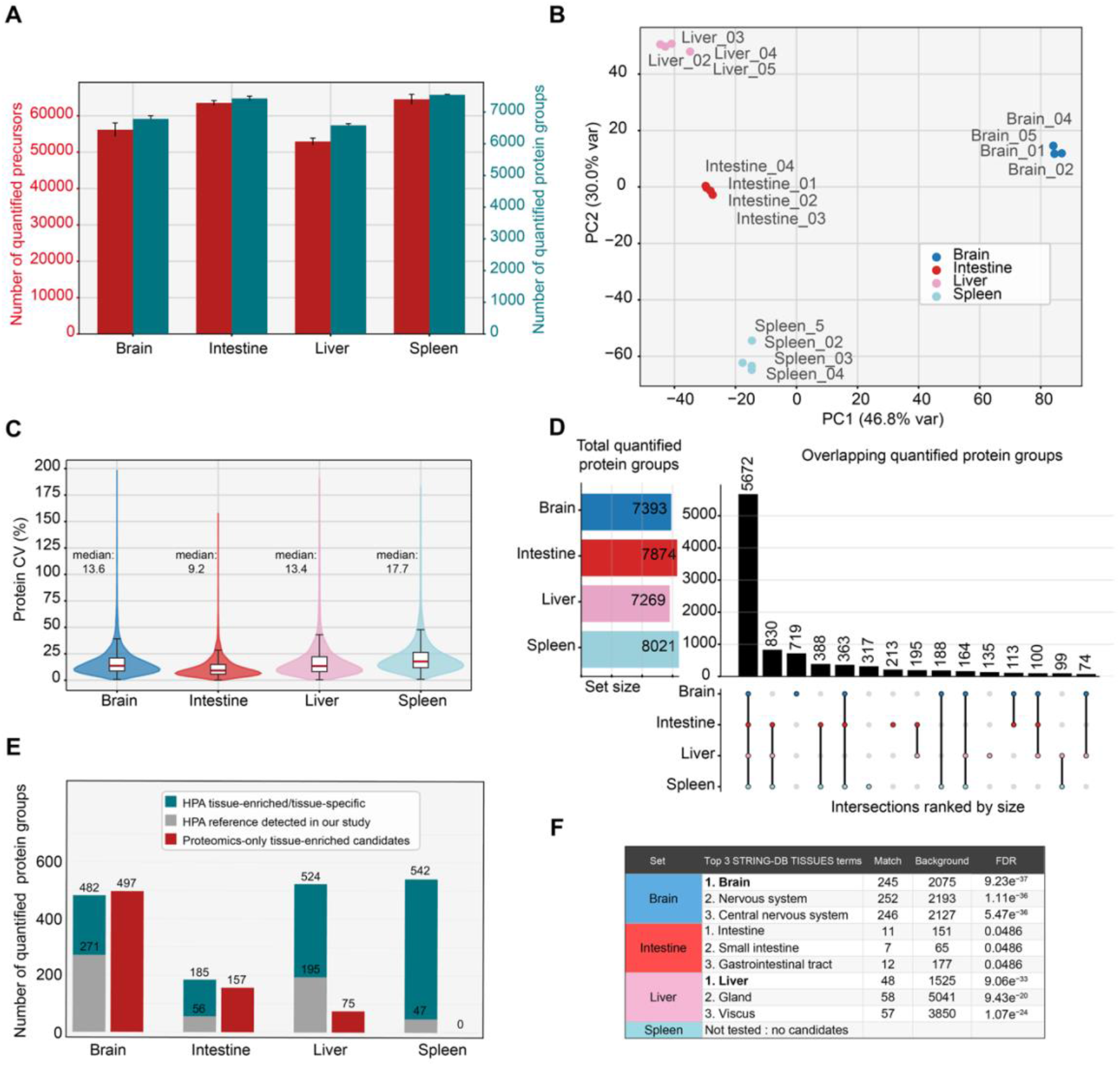
Cross-tissue validation of the LCM-based spatial proteomics workflow. **(A)** Proteome coverage obtained from FFPE mouse brain, intestine, liver, and spleen tissue sections. Bar plots show the number of quantified precursors and quantified protein groups per tissue. Error bars indicate the standard deviation across replicate samples. **(B)** Principal component analysis (PCA) of protein-group intensities showing separation of samples according to tissue type. **(C)** Distribution of protein-level coefficients of variation (CVs) across replicate samples for each tissue. Violin plots with overlaid boxplots are shown, and median CV values are indicated above each group. **(D)** Distribution of quantified protein groups across tissues. Left: total number of quantified protein groups per tissue. Right: UpSet plot showing shared and tissue-specific quantified protein groups. **(E)** Comparison of tissue-enriched protein groups defined in this study with tissue-enriched or tissue-specific annotations from the Human Protein Atlas. Bars indicate HPA tissue-enriched/tissue-specific protein groups, HPA reference protein groups detected in our dataset, and proteomics-only tissue-enriched candidates. **(F)** STRING-DB TISSUES enrichment analysis of tissue-enriched protein groups not annotated as tissue-enriched in the Human Protein Atlas, showing enriched tissue-related terms for each organ.

### A spatial proteomic atlas of the mouse brain choroid plexus

To demonstrate the versatility of our workflow in a complex biological discovery setting, we applied the pipeline to characterize the proteomic landscape of the mouse brain choroid plexus across multiple biological variables, including sex and age and different regions specifically the lateral ventricular and central (third ventricle) plexuses.

The choroid plexus (ChP) is a specialized and highly vascularized structure located within the ventricular system of the brain, including the lateral, third, and fourth ventricles. It plays a fundamental role in maintaining central nervous system (CNS) homeostasis by producing cerebrospinal fluid (CSF) and regulating the exchange of molecules between the blood and the brain environment.

Despite the increasing recognition of its physiological and pathological importance, the molecular composition of the choroid plexus remains incompletely characterized. Most large-scale studies have focused on transcriptomic profiling, including bulk RNA sequencing and single-cell RNA sequencing approaches that have revealed cellular heterogeneity and transcriptional programs within the ChP epithelium and stromal compartments^25,26^. However, comprehensive proteomic datasets of the choroid plexus remain limited.

In this study, we utilized a cohort of 12 mice, balanced by sex (6 male, 6 female) and age (8–10 weeks vs. 41–47 weeks). We specifically targeted three regions: the lateral choroid plexus (right and left) and the central choroid plexus, accessible from the same slice. The integration of biological triplicates and technical duplicates ensured high quantitative rigor for the final dataset and resulted in the analysis of 72 distinct anatomical regions (Figure 4A). Across the entire cohort, the pipeline achieved deep proteomic coverage, identifying on average 7,000 protein groups and 62,000 identified precursors per run (Figure 4B), resulting in 8483 identified protein groups across the 72 runs.

**Figure 4.**
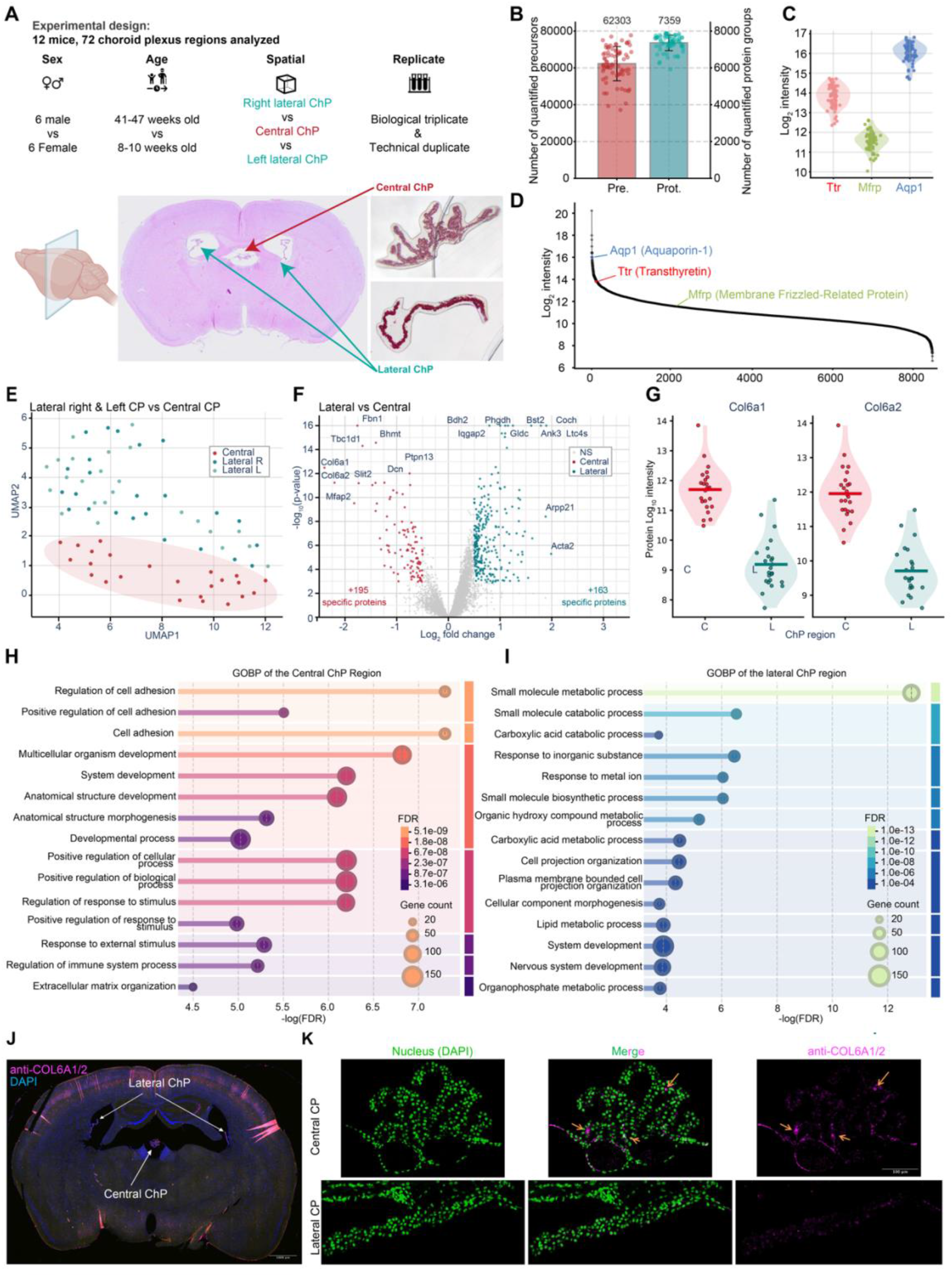
Spatial proteomic profiling of the mouse choroid plexus. **(A)** Experimental design. Choroid plexus regions were isolated from 12 mice balanced by sex and age, including right lateral, left lateral, and central choroid plexus regions, with biological triplicates and technical duplicates. Representative histological images illustrate the sampled regions. **(B)** Proteome coverage across choroid plexus samples. Bar plots show the mean number of quantified precursors (Pre.) and quantified protein groups (Prot.) per run. Error bars indicate the standard deviation across samples. **(C)** Relative abundance of representative choroid plexus marker proteins, including Ttr, Mfrp, and Aqp1. **(D)** Ranked protein-group intensity distribution highlighting representative choroid plexus marker proteins, including Aqp1, Ttr, and Mfrp. **(E)** UMAP projection of choroid plexus samples showing separation between central and lateral regions (left and right). **(F)** Volcano plot comparing lateral (green) and central (red) choroid plexus regions, highlighting representative region-enriched protein groups. **(G)** Relative abundance of Col6a1 and Col6a2 in central and lateral choroid plexus regions. **(H)** Gene Ontology Biological Process enrichment analysis of protein groups enriched in the central choroid plexus. **(I)** Gene Ontology Biological Process enrichment analysis of protein groups enriched in the lateral choroid plexus. **(J)** Immunofluorescence overview of COL6A1/2 staining (pink) in a coronal mouse brain section; DAPI labels nuclei (blue). **(K)** Representative higher-magnification images of central and lateral choroid plexus showing DAPI (green), merged signal, and anti-COL6A1/2 staining (pink). Arrows indicate accumulation of COL6A1/2 signal in the central choroid plexus compared with lateral regions.

Characteristic choroid plexus markers, including transthyretin (TTR)^27^ and aquaporin-1 (AQP1)^28^, together with membrane frizzled-related protein (MFRP), an epithelial membrane protein reported as tissues specific in the choroid plexus in the Human Protein Atlas^29^, were consistently detected and ranked among the most abundant proteins, confirming accurate spatial sampling of this structure (Figure 4C–D). The reproducible detection of these markers, particularly membrane-bound and secreted proteins, supports the suitability of the combined DDM lysis and SDB-RPS cleanup workflow for proteomic analysis of the lipid-rich choroid plexus. Unsupervised clustering via Uniform Manifold Approximation and Projection (UMAP) revealed that spatial orientation, central versus lateral, was the dominant factor influencing the proteomic profile (Figure 4E). The central ChP formed a distinct cluster separated from both the left and right lateral ChP regions. The two lateral regions showed substantial overlap, suggesting a relatively symmetrical molecular identity. While demographic variables such as sex and age contributed to the overall variation, their impact was secondary to anatomical compartmentalization of the tissue (Figure S4A–D).

Differential abundance analysis identified robust molecular signatures, distinguishing the central and lateral ChP compartments. The central ChP was strongly enriched in extracellular matrix and structural proteins, including FBN1, COL6A1, COL6A2, DCN, and MFAP2 (Figure 4F). Additional central-enriched proteins, such as SLIT2, WFIKKN2, THSD7A, FN1, VWF, CTHRC1, and AKAP12, further support a profile dominated by matrix organization, cell–matrix interactions, vascular-associated remodeling, and barrier-supportive architecture. This suggests that the central ChP may possess a more pronounced structural and stromal specialization, potentially reflecting differences in tissue anchoring, vascular organization, and epithelial barrier maintenance.

Conversely, the lateral ChP was enriched in proteins associated with metabolic specialization, ion handling, cellular stress responses, and neural-interface functions. Among the lateral-enriched proteins were BDH2 and PHGDH, enzymes linked to ketone-body-related metabolism and serine biosynthesis, respectively. Additional lateral-enriched proteins included COCH, LTC4S, DPEP1, CA4, BST2, HKDC1, CBS, and MT1, suggesting increased capacity for small-molecule metabolism, redox regulation, immune-related signaling, and pH/ion homeostasis. The enrichment of ACTA2, TAGLN, and VCAN may also indicate regional differences in vascular or perivascular components within the lateral ChP. Together, these data suggest that lateral ChP regions are more metabolically and signaling-active, consistent with their interface with ventricular CSF circulation.

Gene Ontology Biological Process (GOBP) enrichment analysis corroborated these spatial trends. Protein groups enriched in the central ChP were associated with cell adhesion, extracellular matrix organization, anatomical structure morphogenesis, and regulation of immune system processes (Figure 4H), consistent with a more structural and matrix-rich phenotype. In contrast, protein groups elevated in the lateral ChP were overrepresented in categories including small molecule metabolic processes, carboxylic acid catabolic processes, organic hydroxy compound metabolism, and nervous system development (Figure 4I). These findings suggest enrichment of proteins involved in metabolite processing and small-molecule metabolism within the lateral ChP, potentially reflecting regional specialization associated with its interface with ventricular CSF. Because COL6A1 and COL6A2 were among the most strongly enriched proteins in the central ChP (Figure 4F-G), we validated their spatial distribution by immunofluorescence. Consistent with the proteomic analysis, COL6A1/2 immunoreactivity was abundant throughout the central ChP but substantially reduced in the lateral ChP (Figure 4J-K), supporting a regionally specialized extracellular matrix-rich phenotype in the central ChP. Overall, these findings reveal clear spatial proteomic compartmentalization of the choroid plexus, with the central region characterized by extracellular matrix and structural organization, and the lateral regions characterized by metabolic, transport, and signaling-related functions.

To complement the GO enrichment analysis, a Shared and Unique Structures (SUS) plot analysis was used to visualize the contribution of individual proteins to the regional separation and to group them into broad biological modules. This analysis confirmed that the main proteomic difference occurred between central and lateral choroid plexus compartments, with only limited separation between the left and right lateral regions (Figure S5). It also confirmed that proteins enriched in the central compartment were predominantly associated with extracellular matrix and tissue remodeling, vascular/perivascular organization, barrier-related functions, and reactive or inflammatory signatures. In contrast, lateral-enriched proteins were distributed across metabolic/mitochondrial, neuronal/synaptic, myelin/oligodendrocyte-associated, and contractile/perivascular modules (Figure S6). Together, these module-level patterns support the UMAP and GO enrichment analyses and indicate that the choroid plexus proteome is primarily organized along a central–lateral axis, while the left and right lateral regions share a largely overlapping molecular identity.

To determine whether biological sex or age influenced the choroid plexus proteome, we compared male and female mice and contrasted young (8–10 weeks) versus old (41–47 weeks) animals within each spatial compartment. Sex accounted for a much smaller proportion of variance than spatial location, although a subset of proteins exhibited significant sexual dimorphism. Male choroid plexus samples displayed higher abundance of several major urinary proteins, including MUP1, MUP2, MUP17, and MUP20, as well as the serine protease inhibitor SERPINA3K and the apolipoproteins APOA1 and APOA2. In contrast, females showed elevated levels of DEAD-box helicase 3 X-linked (DDX3X) and a small set of other female-biased proteins, including GLRX1. These differences did not overlap substantially with the region-specific markers identified above, indicating that sex-associated proteomic variation was largely independent of spatial compartmentalization (Figure S4A–B). Comparison of old and young mice revealed a more pronounced age-dependent proteomic signature. Old animals displayed increased abundance of immunoglobulin-related proteins, including IGKC, IGHG2B, IGHM, and IGKV12-41, together with immune and inflammatory proteins such as FCGR2, MPEG1, MNDAL, VAV1, CD180, CASP4, CRP, SAA4, HP, C8A, C9, and MASP2. Additional proteins highlighted in the volcano plot, including HTRA1, ADAMTSL4, and ASS1, suggested age-associated extracellular matrix remodeling and tissue adaptation. In contrast, young animals were enriched in proteins linked to tissue structure and development, including FREM2, FRAS1, LAMA1, CENPA, TPBG, and NEGR1 (Figure S4C–D). Gene Ontology enrichment analysis of proteins elevated in old animals further supported these observations, identifying significant enrichment for extracellular matrix organization, collagen fibril organization, regulation of cell adhesion, acute inflammatory response, lymphocyte-mediated immunity, and adaptive immune response pathways (Figure S4E). Together, these findings indicate that ageing of the choroid plexus is accompanied by increased immune-related and extracellular matrix remodeling signatures, alongside reduced representation of developmental and structural proteins.

Overall, these results demonstrate that the choroid plexus exhibits subtle but detectable proteomic signatures of sex and age that are superimposed on the dominant spatial specialization described above. To permit a more detailed exploration of the findings, an associated website is under development and will go live in the upcoming weeks.

## Discussion

Over the last five years, biomedical research has entered the era of spatial biology. Recognition of the importance of preserving spatial context when analyzing molecular profiles, together with major technological advances, has propelled the field and transformed how scientists visualize and dissect disease mechanisms. Several milestone studies in LCM MS-based spatial proteomics have provided invaluable methodological insights^6,30,31^. In this context, our study highlights the value of combining complementary technological platforms and demonstrates that high-quality spatial proteomic data can be generated without necessarily relying on the latest instrumentation.

Maximizing the quality of the final output required careful, stepwise optimization of the entire workflow. Starting with histology, different tissue types and preservation methods required specific adjustments to slide pre-processing and section thickness. On the imaging side, integration with OMERO and QuPath for image-based region selection, segmentation, and annotation enabled precise microdissection guided by spatial cellular features. For sample preparation, a simple one-pot reaction combined with standard desalting generated high-quality samples. The mixed-mode retention mechanism of SDB-RPS enabled efficient removal of detergents and other contaminants through high-organic washes, followed by peptide elution through a pH shift, yielding clean peptide mixtures that supported robust quantification. Careful optimization of mass spectrometry acquisition on a mainstream and relatively accessible platform produced identification and quantification results comparable to those obtained with more recent high-end instruments. These findings suggest that existing instrumentation can remain highly valuable when paired with optimized workflows, which is particularly relevant in the context of current funding constraints.

The developed workflow naturally entails certain constraints. The main limitation is the moderate throughput of the pipeline, driven by semi-manual handling at the microscope and the length of each MS acquisition. Nevertheless, we have successfully applied it across multiple projects, including studies involving up to 100 samples. Furthermore, the method can be readily adapted to different Orbitrap-based instruments and to shorter gradients on more sensitive systems without necessarily compromising analytical depth or quantitative performance. An additional limitation is that although the workflow provides substantial proteome depth from as few as 20 cells, it does not generate single-cell proteomic data. However, by pooling individual cells or microregions of a defined phenotype, the approach provides sufficient analytical depth, reduces inter-sample variability, and still generates biologically meaningful spatial information.

The analysis of the choroid plexus not only showcased the advantages of our pipeline but also provided insight into the functional organization of this tissue. Spatial proteomic profiling of the choroid plexus (LL vs LR vs 3C) revealed a clear separation between central and lateral ventricular regions, while the left and right lateral regions showed little separation from one another, indicating a largely shared lateral molecular identity. Unsupervised clustering and differential abundance analysis identified region-specific protein signatures distinguishing these compartments. Functional enrichment analysis indicated that proteins elevated in the lateral choroid plexus were predominantly associated with metabolic and small-molecule processing pathways, whereas proteins enriched in the central choroid plexus were linked to cell adhesion, extracellular matrix organization, anatomical structure morphogenesis, and immune-related processes. These observations are consistent with ventricle-specific transcriptional programs previously reported in single-cell transcriptomic studies of the choroid plexus, suggesting that regional molecular specialization is preserved at the protein level^26^.

Interestingly, previous studies have reported sex-related differences in choroid plexus size or volume, potentially reflecting hormonal regulation or sex-dependent ventricular physiology^32,33^. In our dataset, however, sex accounted for only a minor fraction of proteomic variation compared with anatomical location. Similarly, the left and right lateral choroid plexus showed little separation at the proteomic level, suggesting that potential morphological or physiological asymmetries are not accompanied by major differences in protein-level functional specialization in this dataset. Age-associated remodeling of the choroid plexus has previously been linked to increased inflammatory signaling and altered immune regulation within the ventricular system. Transcriptomic studies have reported elevated interferon and cytokine signaling pathways in the aging choroid plexus, supporting the idea that this structure acts as a key interface between peripheral immune signals and the brain^26,34,35^. In our proteomic dataset, aging was associated with increased abundance of immune-and inflammation-related proteins, while young animals showed higher levels of proteins linked to tissue structure and developmental organization. These age-associated protein signatures may therefore represent candidate markers for monitoring choroid plexus remodeling in disease contexts, particularly in conditions involving neuroinflammation, blood–CSF barrier dysfunction, altered CSF composition, or impaired immune surveillance. This is especially relevant given that aging-related inflammatory activation of the choroid plexus has been linked to impaired brain function^34^, and that ventricle-specific choroid plexus programs are altered in disease-relevant inflammatory contexts^26^. These age-associated protein signatures may therefore represent candidate markers for monitoring choroid plexus remodeling in disease contexts involving neuroinflammation, blood–CSF barrier dysfunction, altered CSF composition, or impaired immune surveillance.

In the near future, we plan to expand our workflow toward molecular-guided microdissection of tissue regions, in which areas of interest are selected based on their distinct molecular signatures rather than solely on histology. For example, this could involve integrating complementary spatial readouts such as MALDI imaging-based lipidomics^36^ and/or multiplex immunofluorescence^4^ acquired on the same slide to generate maps that highlight specific lesion subregions which can be used to define for dissection. In addition, we are interested in extending the pipeline to signaling-focused studies by adapting it for phosphoproteomics. This will require tailoring sample handling and processing to preserve labile phosphorylation events and implementation of enrichment strategies compatible with low-input material. Finally, expansion microscopy^37^ could become a valuable addition for projects that demand subcellular resolution. By physically expanding the specimen prior to imaging, this approach can facilitate more precise localization of structures and markers and enable microdissection strategies that better reflect subcellular organization. Together, these developments aim to broaden the range of biological questions that can be addressed by our platform.

## Methods

### Animals

C57Bl6/J mice were kept in a Specific Pathogen Free (SPF) environment with a 12h light and 12h dark cycle and *ad libitum* access to food and water. For animal experimentation, all procedures were performed according to protocols approved by the Veterinary Authorities of the Canton Vaud and according to the Swiss Law (License VD 3615, EPFL) and were in accordance with the ARRIVE guidelines and 3R principle (Reduction, Replacement, Refinement) for laboratory animals.

| ID | Age | Sex |
| --- | --- | --- |
| 19253 | Young / 8w | female |
| 19254 | Young / 8w | female |
| 19268 | Young / 9w | female |
| 19215 | Young / 10w | male |
| 19252 | Young / 8w | male |
| 19266 | Young / 9w | male |
| 18387 | Old / 47w | female |
| 18195 | Old / 47w | female |
| 18196 | Old / 47w | female |
| 17955 | Old / 47w | male |
| 18422 | Old / 41w | male |
| 18423 | Old / 41w | male |

### Histological Preparation

Mouse brain tissues were collected and fixed in 4% paraformaldehyde for 24 h, then processed and embedded in paraffin. The resulting FFPE blocks were stored at 4 °C until sectioning. For slide preparation, blocks were sectioned at 4 µm thickness, and sections were mounted alternately on SuperFrost® Plus slides (#J1800AMNZ) for imaging or on Leica 2.0 µm PEN membrane slides (#11505189) for microdissection.

Prior to downstream processing, mounted sections were deparaffinized using standard histological procedures. This included heat-assisted softening of the paraffin, organic solvent-based removal with xylene, and graded rehydration through decreasing concentrations of ethanol (96%, 80%, and 70%). Sections destined for microdissection were lightly stained with hematoxylin and eosin to aid structural recognition during ROI selection. After staining, slides were air-dried at room temperature and stored at 4 °C until laser microdissection.

### Image Acquisition

Histological slides were imaged, in brightfield, with a 20x/0.80 air objective, on a VS200 slide scanner microscope from Evident. Once acquired, images were uploaded to the available OMERO database.

### Drawing of ROIs

QuPath analysis software, v0.6.0, was used to precisely draw ROIs. Images were imported in a QuPath project, through the QuPath-OMERO extension v0.2.2. ROIs were manually drawn around the tissue, taking care to only draw closed-shape ROIs, with or without holes. Different classes were assigned to the ROIs according to what they represented. To include the cutting laser thickness, ROIs were expanded by 1um distance all around their border, using a home-made groovy script (https://github.com/BIOP/omero-lmd), runnable directly within QuPath. The well ID, corresponding to the tube into which the cut will fall during microdissection, is also assigned to each ROIs. Three points were precisely positioned on calibrated slide patterns and were used as calibration points for microdissection. Each point was named with index-based names, to recognize each point independently. ROIs and well ID were finally sent to OMERO database and linked to the corresponding image in the form of a csv file.

### Laser Microdissection

For preliminary pipeline developments and fresh frozen tissue experiments (Figure S3), samples were laser microdissected using Zeiss PALM Microbeam Laser Microdissection microscope equipped with a SingleTube 500 CM II tube holder and controlled by PALMRobo 4.6 Pro software. Dissections were performed in brightfield mode using a Zeiss Plan-Neofluar 20x/0.5 air objective. The following laser settings were used: cutting speed at 20%, cutting energy at 60%, cutting focus at 75%, LPC energy at 81%, and LPC focus at 80%. Tissue samples were catapulted into 0.2 mL opaque adhesive cap tubes. The CapCheck function was employed to ensure efficient tissue recovery.

Due to the capacity to receive annotation and be linked to OMERO the Leica laser cutting microscope was favored for further development. For all FFPE experiments and choroid tissues, a LMD6500 (Leica Microsystems) laser microdissection microscope was used to cut ROIs retrieved from the OMERO database. A home-made python script (https://github.com/BIOP/omero-lmd), based on py-lmd library^38^, was run to convert shapes from OMERO format to LMD-readable format, including the well ID where the cut will fall into. Due to a limitation of the LMD 8.4 software, which doesn’t allow a shape contour to be described with more than 500 points, a shape simplification was performed during the conversion with Ramer–Douglas–Peucker algorithm (https://shapely.readthedocs.io/en/2.1.2/reference/shapely.simplify.html) The xml file resulting from the conversion was then imported in the LMD 8.4 software. The following laser settings were used: using Final Pulse mode: Power: 30%, Aperture: 2, speed: 10. Tissue samples were isolated into 0.2 mL opaque cap tubes. The collected Choroid plexus tissue regions ranged from 80 000 µm^2^ to 450 000 µm^2^. Each region was manually monitored under a binocular to carefully assess the isolation.

### Sample Preparation for LC-MS Analysis

Under a 20× binocular microscope, tissue samples were carefully resuspended from the caps and transferred into 0.2 mL Eppendorf tubes using 100 µL of acetonitrile. Samples were pelleted and dried by vacuum centrifugation.

Dried tissue samples were resuspended in 15 µL of decrosslinking buffer containing 300 mM Tris-HCl, pH 8.0, and 0.2% DDM. FFPE samples were incubated at 80 °C for 90 min, followed by progressive cooling to 37 °C. Digestion was initiated by adding 15 µL of a concentrated digestion mixture prepared from 50 µL of 100 mM tris(2-carboxyethyl)phosphine (TCEP), 50 µL of 400 mM chloroacetimide (CAA), 50 µL of 1% DDM, 10 µL of 100 ng/µL trypsin, 10 µL of 100 ng/µL Lys-C, and water to a final volume of 250 µL. After addition to the sample, the final reaction contained 150 mM Tris-HCl, pH 8.0, 10 mM TCEP, 40 mM CAA, 0.2% DDM, and 2 ng/µL each of trypsin and Lys-C in a final volume of 30 µL. Samples were digested at 37 °C overnight.

Resulting peptides were desalted on SDB-RPS StageTips^14^ dried by vacuum centrifugation and stored at −20 °C until LC–MS analysis.

### LC-MS Methods

Peptides were reconstituted in 20 µL of 2% acetonitrile and 0.1% formic acid in Milli-Q water. The solution was agitated at 800 rpm for 20 min at room temperature using a Thermomixer Compact (Eppendorf) and then briefly centrifuged. Peptides were injected and separated into a VANQUISH HPLC (ThermoFisher Scientific, USA) connected online to an Exploris™ 480 Orbitrap mass spectrometer (ThermoFisher Scientific, USA). A capillary precolumn (Acclaim™ PepMap™ C18; 3 µm, 100 Å; 2 cm × 75 µm ID; ThermoFisher Scientific, USA) was used for sample trapping and cleaning. Analytical separations were performed at 250 nL/min over a 150 min biphasic gradient on a 50 cm in-house–packed capillary column (75 µm ID; ReproSil-Pur C18-AQ; 1.9 µm silica beads; Dr. Maisch, Ammerbuch, Germany).

Data were recorded in data-independent acquisition (DIA) mode using SM-DIA method described below.

### MS method development

All DIA method development was assessed using the Pierce HeLa Protein Digest Standard (Thermo Fisher Scientific, 88328). The lyophilized protein digest (20 μg) was reconstituted in 200 μL of 2% acetonitrile (ACN) in 0.1% formic acid (FA) and mixed thoroughly at room temperature for 10 min to obtain a final concentration of 100 ng/μL. This stock solution was further diluted 20-fold and stored at −20 °C until use.

To identify the most prominent peptides across a broad mass range, 300 ng of HeLa peptides were analyzed using data-independent acquisition (DIA) with a wide full MS scan range (m/z 350–1650), as previously described^19^. Full MS acquisition parameters included a resolution of 120,000, a maximum ion injection time of 20 ms, and an automatic gain control (AGC) target of 300%. Each MS survey scan was followed by DIA MS/MS scans acquired at a resolution of 30,000, with a maximum ion injection time of 55 ms, an AGC target of 3000%, stepped normalized collision energies of 25.5%, 27%, and 30%, and a fixed first mass of m/z 200.

Subsequently, different DIA acquisition strategies were evaluated, as described in the results section and summarized in Supplementary Data 1. MS1 scans were acquired at a resolution of 60,000, with an AGC target of 300% and the maximum injection time set to auto. Fixed-width DIA windows, with total *m/z* spans corresponding to multiple of 14 Da (ranging from 12 to 40 × 14 Da), were scanned across the 350–1650 *m/z* range in 2 Da increments. For each window size configuration, the window position capturing the largest number of precursor ions was identified. To assess the impact of progressively decreasing precursor-range widths on DIA performance, four total isolation ranges (560, 392, 252, and 196 Da) were selected. For MS2 DIA scans, isolation windows were distributed to maintain an equal number of precursor ions per window. Resolution was included as a variable parameter in the method evaluation (see Supplementary Data 1). The maximum injection time was set to auto, and stepped normalized collision energies of 22%, 26%, and 30% were applied. The MS2 AGC target was set to 3000%, with a fixed scan range of m/z 200–1800.

For the assessment of the spatial proteomics workflow and the brain choroid plexus study, the best-performing DIA method was selected. Briefly, MS1 scans were acquired using a scan range between 418-670 m/z using a resolution of 60,000, with an automatic gain control (AGC) target of 300% and the maximum injection time set to auto. MS2 isolation windows were distributed to maintain an equal number of precursor ions per window. The resolution was set to 120,000 (at m/z 200), and the maximum injection time was set to auto. Stepped normalized collision energies of 22%, 26%, and 30% were applied. The MS2 AGC target was set to 3000%, with a fixed scan range of m/z 200–1800.

A three-proteome benchmark sample was prepared with expected log₂ fold changes (A/B) of 0 for human, +1 for yeast, and −2 for *E. coli*, using Pierce HeLa Protein Digest Standard (Thermo Fisher Scientific, 88328), MS-Compatible Yeast Protein Digest (Promega, V7461), and MassPREP *E. coli* Digest Standard (Waters, 186003196). Two mixtures were generated with defined ratios: Sample A contained 72% human, 24% yeast, and 4% *E. coli*, while Sample B contained 72% human, 12% yeast, and 16% *E. coli*. Both samples were freshly diluted to 5 ng/μL (total protein), and injection of 2 μL of either sample resulted in a total load of 10 ng on column.

### Data Analysis

#### MS-method development

Raw data were analyzed with DIA-NN (v2.1.0)^39^ using library free search against the *Homo sapiens* SwissProt database (20’405 sequences, Release 2025_04, complemented with the common MaxQuant contaminants^40^. Results were filtered at 1% FDR. Search parameters included: minimum fragment m/z at 200, maximum fragment m/z at 1800, N-terminal methionine excision enabled, in silico digest with cuts at K* and R*, maximum number of missed cleavages set to 1, minimum peptide length set to 7, maximum peptide length set to 30, minimum precursor m/z at 300, maximum precursor m/z at 1800, minimum precursor charge set to 1, maximum precursor charge set to 4. Carbamidomethylation (C) was set as a fixed modification, whereas oxidation (M) as a variable modification. Maximum number of variable modifications was set to 2 and Match between runs (MBR) was enabled. The Unrelated runs option was checked. The resulting DIA-NN parquet file was read with in-house generated R programs to draw graphs. Precursors were filtered with Lib.Q.Value <= 0.01 & Q.Value <= 0.01 and protein groups with Lib.PG.Q.Value <= 0.01.

#### Triple Proteome

Raw data were processed using DIA-NN (v2.1.0) against a combined database comprising the *Homo sapiens* UniProt proteome (20,405 sequences; release 2025_04), the *Saccharomyces cerevisiae* UniProt proteome (6,066 sequences; release 2025_04), and the *Escherichia coli* UniProt proteome (4,402 sequences; release 2025_04), supplemented with common MaxQuant contaminants. Data were processed using the same parameters as in the method development study.

The resulting DIA-NN report.*pg_matrix* output was further processed to remove potential contaminants. Protein groups were filtered to retain only those supported by at least one proteotypic peptide. Protein groups containing at least 75% of valid values in at least one group were conserved for further analysis. No imputation on the maxLFQ intensities was performed.

#### FFPE and Fresh frozen mouse tissues analysis

Raw data were processed as described above for the triple-proteome analysis, using DIA-NN (v2.1.0). Data were searched against the *Mus musculus* Swiss-Prot proteome (17,237 sequences; release 2025_03), supplemented with common MaxQuant contaminants. The resulting DIA-NN *pg_report* files were processed as described above, including removal of potential contaminants and retention of protein groups supported by at least one proteotypic peptide. Data processing and statistical analyses were performed in Python (v3.12) using NumPy (v2.3.3), pandas, SciPy (v1.16.2), and Matplotlib (v3.10.6). For extended analyses (PCA, TAU score), missing values were imputed using a Perseus-like algorithm in Python.

### Choroid plexus

Raw data were analyzed using DIA-NN (v2.1.0) with the same options as the search for the Method study against the *Mus musculus* UniProt database (17’237 sequences, Release 2025-03), complemented with the common MaxQuant contaminants [DOI: 10.1038/nbt.1511]. Minimum and maximum precursor m/z were set respectively at 400 and 800. Resulting DIA-NN reports were converted to MSstats format using MSstatsConvert (v1.18.1) and analyzed in R (v.4.5.1) with MSstats^41^ (v4.16.1). Differential protein expression analysis was performed using the linear mixed-effects model accounting for paired samples, followed by the Benjamini–Hochberg procedure. Adjusted P value lower than 0.01 (and |log2 fold change| > 0.5) were considered as significant.

### Gene Ontology Biological Process enrichment

Protein groups with FDR < 0.01 from the central versus lateral choroid plexus comparison were submitted to STRING DB for Gene Ontology Biological Process enrichment analysis without applying an additional log₂ fold-change cutoff. The background was defined as all protein groups quantified across the choroid plexus dataset. GO terms were clustered using a similarity threshold of 0.5, and significantly enriched terms were filtered according to the STRING enrichment FDR^42^

Prior to multivariate analysis, protein log2-transformed intensities, from MSstats analysis, were filtered to retain proteins with at least 75% valid values in at least one group. Missing values were imputed using a Perseus-like approach (downshift = 1.8SD, width = 0.3SD).

Two Orthogonal Partial Least Squares Discriminant Analysis (OPLS-DA) models were built using the ropls R package (v1.42.0): one contrasting LR versus C, and one contrasting LL versus C. Each model used one predictive component with the number of orthogonal components determined automatically by cross-validation (7-fold). Pareto scaling was applied prior to modeling. Model significance was assessed by 100-permutation testing (pR2Y and pQ2 < 0.01 considered significant).

For each model, the protein p(corr) was computed as the Pearson correlation between its abundance and the predictive score, and the VIP (Variable Importance in Projection) score was extracted using getVipVn method^43^. Proteins were classified on Shared and Unique Structures plot^44^ (SUS-plot) according to their VIP scores (threshold: VIP > 1.5) in each model.

Over-representation analysis (ORA) of Gene Ontology Biological Process (GO-BP) terms was performed for each protein using the method enrichGO from the R clusterProfiler^45^ (v4.18.4) package. (Benjamini–Hochberg adjusted p < 0.01), with all quantified proteins as background. UniProt protein identifiers were converted to Entrez Gene IDs, with org.Mm.eg.db (v3.22.0) as the reference annotation. GO terms with undefined Information Content in org.Mm.eg.db were excluded. Semantic similarity between significant GO terms was computed using the Wang measure of the mgoSim method of the R package GOSemSim^46^ (v2.36.0) and GO terms were clustered by hierarchical clustering (complete linkage) on the resulting distance matrix. Results were visualized as a heatmap of log2(FoldEnrichment) using ComplexHeatmap ComplexHeatmap^47^ (v2.26.1), with rows ordered by semantic similarity clustering (k = 18 clusters).

### ProteomeXchange

The mass spectrometry proteomics data has been deposited to the ProteomeXchange Consortium^48^ via the PRIDE partner repository with the dataset identifier PXD081212.

## Author Contributions

M.P. and T.C. conceived the study, designed the experiments, and wrote the manuscript. T.C performed and/or contributed to all experiments J.SD. designed the study and developed the histology workflow. R.H. developed and optimized the mass spectrometry methodology. F.A. and T.C. performed data analysis and prepared the figures. F.A. designed the website for displaying the results of the Spatial Proteomics Mouse Choroid Plexus Atlas R.D. implemented and integrated the Python–OMERO analysis pipeline. All authors reviewed and approved the final manuscript.

## Acknowledgements

We are grateful to the members of the PCF, in particular Mathilde Willemin, Jonathan Pittet, and Diego Chiappe, for their support and contributions to this study. We also thank the Histology Core Facility, especially Anjalie Schlappi, for the preparation and dissection of mouse brain samples and Gian-Filippo Mancini, Agnes Hautier, Nathalie Muller for their assistance with slide preparation and experimental development. We gratefully acknowledge Florence Mehl for her valuable insights and expert guidance in the multivariate data analysis. We thank Prof. Giovanni D’Angelo for the fruitful discussions and feedback in the creation of the choroid plexus atlas. We are grateful to Dr. Lucie Bracq and Prof. Gisou van der Goot for providing mouse brain samples, which enabled the choroid plexus study.

---

**Figure S1.**
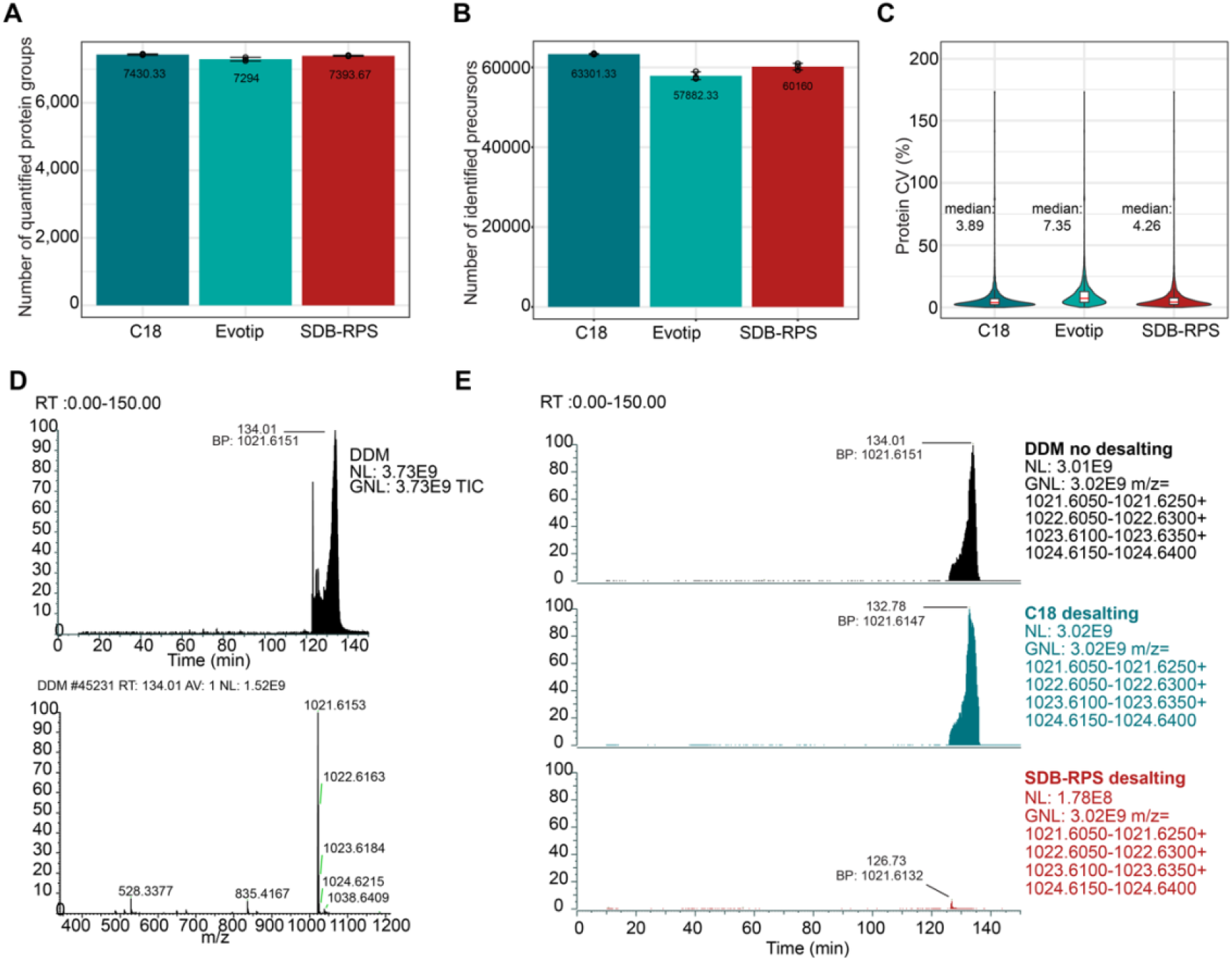
Performance comparison of peptide cleanup strategies for microscale samples. (A, B) Evaluation of peptide recovery using a commercial HeLa peptide mixture desalted with three cleanup formats: C18 & SDB-RPS StageTips (homemade) and manufactured Evotips™ (Evosep). Bar plots show the mean number of quantified protein groups (A) and quantified peptide precursors (B). Error bars indicate the standard deviation across replicates. (C) Distribution of protein-level coefficients of variation (CVs) across cleanup formats. Median protein CV is indicated for each condition. (D) Representative total ion chromatogram (TIC) of a DDM-containing sample showing a high-intensity contaminant signal eluting at late retention time, together with the corresponding MS1 spectrum acquired at RT 134.01 min. The MS1 spectrum shows DDM-related ions, including *m/z* 528.3377 and the dominant isotope cluster around *m/z* 1021.6153–1024.6215. (E) Extracted ion chromatograms of the DDM-related isotope cluster around *m/z* 1021.61–1024.64 in samples processed without desalting, with C18 desalting, or with SDB-RPS desalting. SDB-RPS desalting markedly reduced the DDM-related contaminant signal compared with no desalting and C18 desalting, supporting its selection for detergent-containing low-input samples.

**Figure S2.**
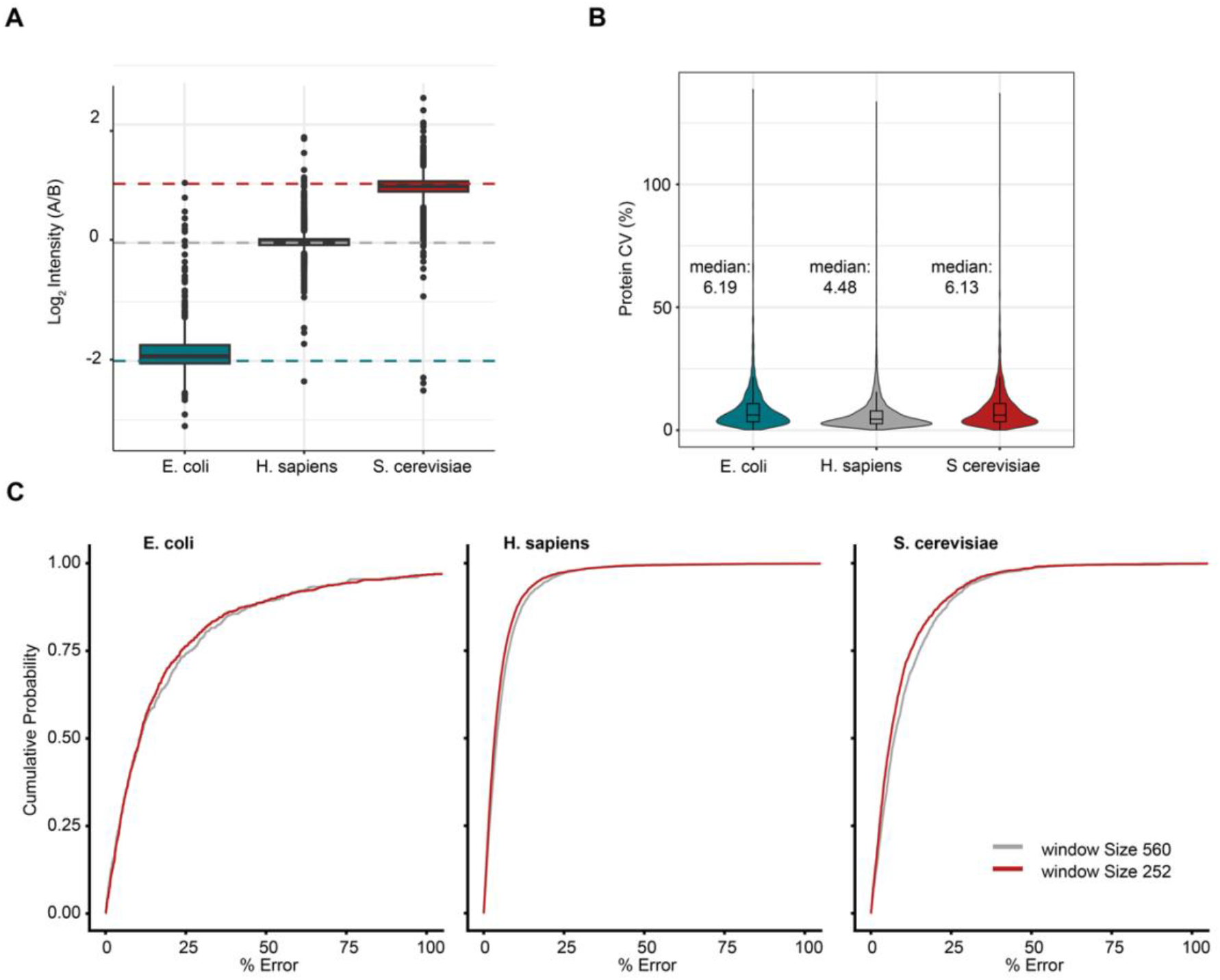
Quantitative accuracy and reproducibility of DIA acquisition methods in a three-proteome benchmark. **(A)** Distribution of measured log₂ fold changes between mixtures A and B for quantified protein groups from *E. coli*, *H. sapiens*, and *S. cerevisiae* using the 256 *m/z* acquisition window. Dashed lines indicate the expected organism-specific fold changes. **(B)** Distribution of protein-level coefficients of variation (CVs) for each organism using the 560 *m/z* acquisition window. Median CV values are indicated above each violin. **(C)** Cumulative distribution functions (CDFs) of protein quantification percent error for each organism, comparing the 252 *m/z* window method in red with the 560 *m/z* window method in grey. Only protein groups with at least 75% valid values in at least one organism group were included.

**Figure S3.**
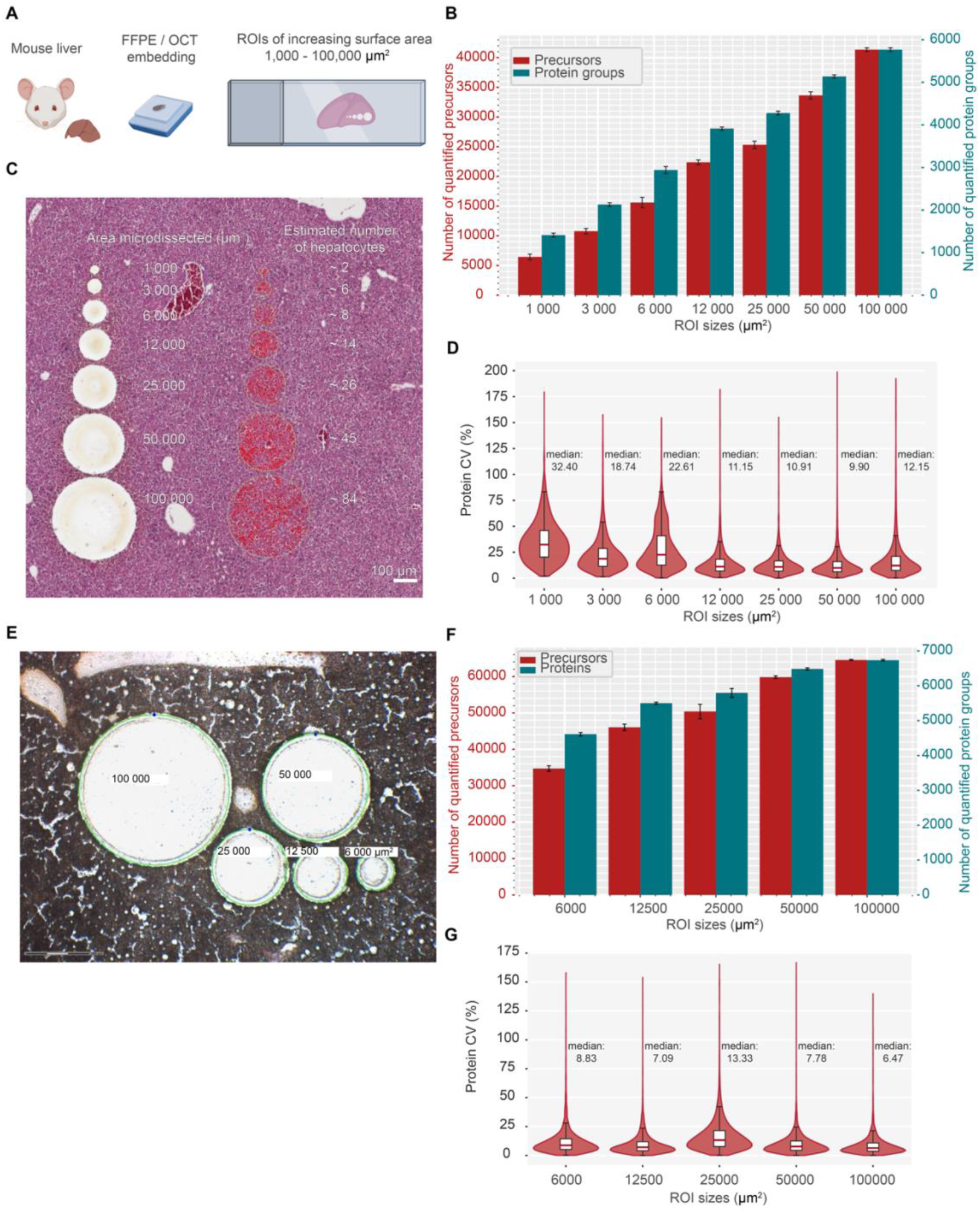
Spatial proteomics performance across ROI sizes in FFPE and fresh-frozen mouse liver tissue. (A) Schematic overview of the ROI-size titration experiment. Mouse liver tissue was prepared as FFPE or fresh-frozen OCT-embedded sections, and circular ROIs of increasing surface area were isolated by laser microdissection. **(B)** Proteome coverage obtained from FFPE liver ROIs of increasing area. Bar plots show the number of quantified precursors (red) and quantified protein groups (green) as a function of ROI size. Error bars indicate the standard deviation across replicate samples. **(C)** Representative histological image of FFPE liver tissue showing circular ROIs of increasing surface area and the corresponding estimated number of hepatocytes determined by QuPath-based cell segmentation. **(D)** Distribution of protein-level coefficients of variation (CVs) across FFPE ROI sizes. Violin plots with overlaid boxplots are shown, and median CV values are indicated above each group. **(E)** Representative histological image of fresh-frozen OCT-embedded liver tissue showing circular ROIs of increasing area. **(F)** Proteome coverage obtained from fresh-frozen OCT liver ROIs of increasing area. Bar plots show the number of quantified precursors and quantified protein groups as a function of ROI size. Error bars indicate the standard deviation across replicate samples. **(G)** Distribution of protein-level coefficients of variation (CVs) across fresh-frozen OCT ROI sizes. Violin plots with overlaid boxplots are shown, and median CV values are indicated above each group

**Figure S4.**
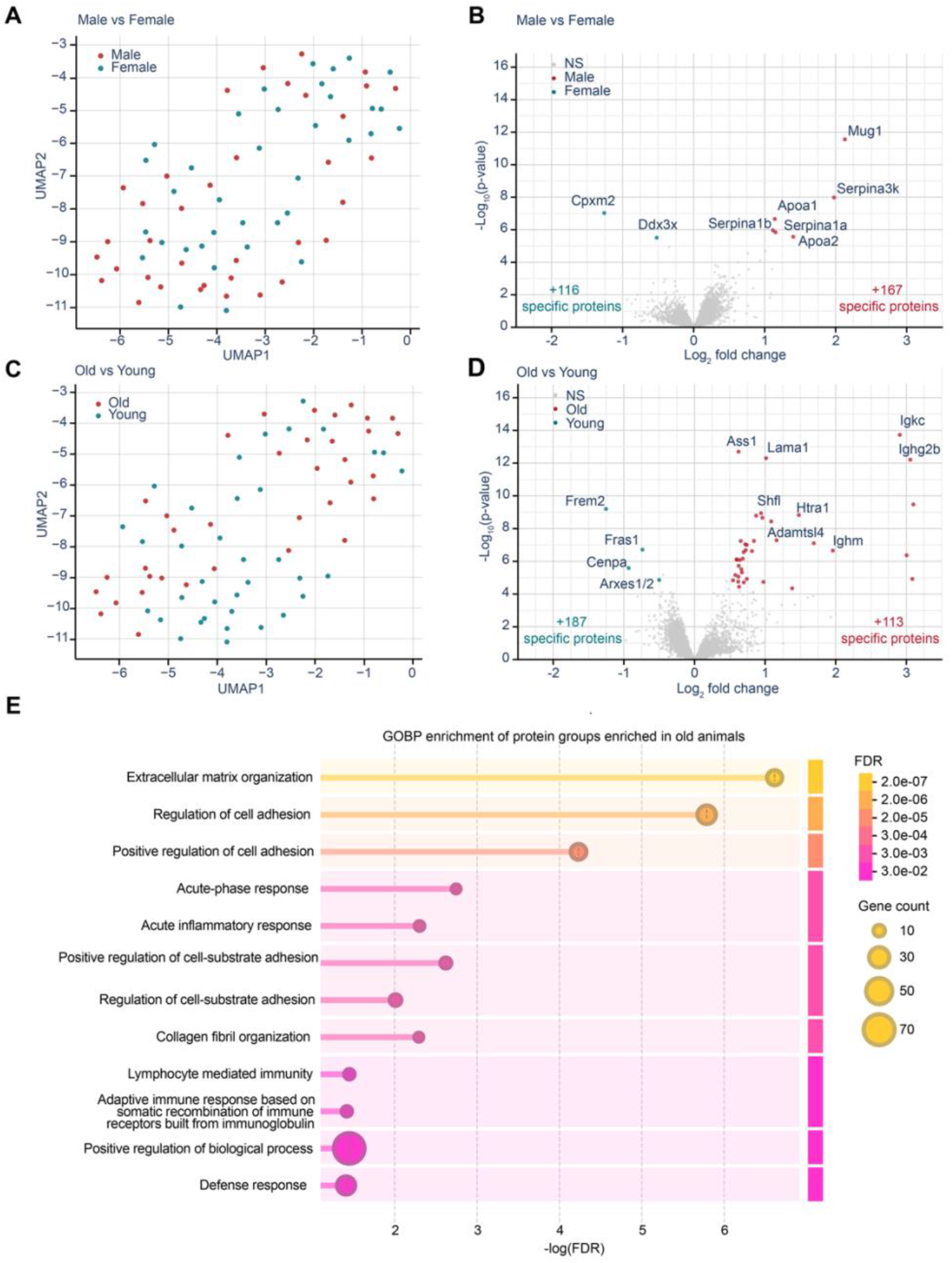
Sex-and age-associated proteomic signatures in the mouse choroid plexus. (A) UMAP projection of choroid plexus samples comparing male and female animals. (B) Volcano plot comparing male and female choroid plexus samples, highlighting representative sex-associated protein groups. Red indicates protein groups enriched in male animals, green indicates protein groups enriched in female animals, and grey indicates non-significant protein groups. (C) UMAP projection of choroid plexus samples comparing old and young animals. (D) Volcano plot comparing old and young choroid plexus samples, highlighting representative age-associated protein groups. Red indicates protein groups enriched in old animals, green indicates protein groups enriched in young animals, and grey indicates non-significant protein groups. (E) Gene Ontology Biological Process enrichment analysis of protein groups enriched in old animals. Dot size indicates gene count, color indicates enrichment FDR, and the x-axis shows −log(FDR).

**Figure S5.**
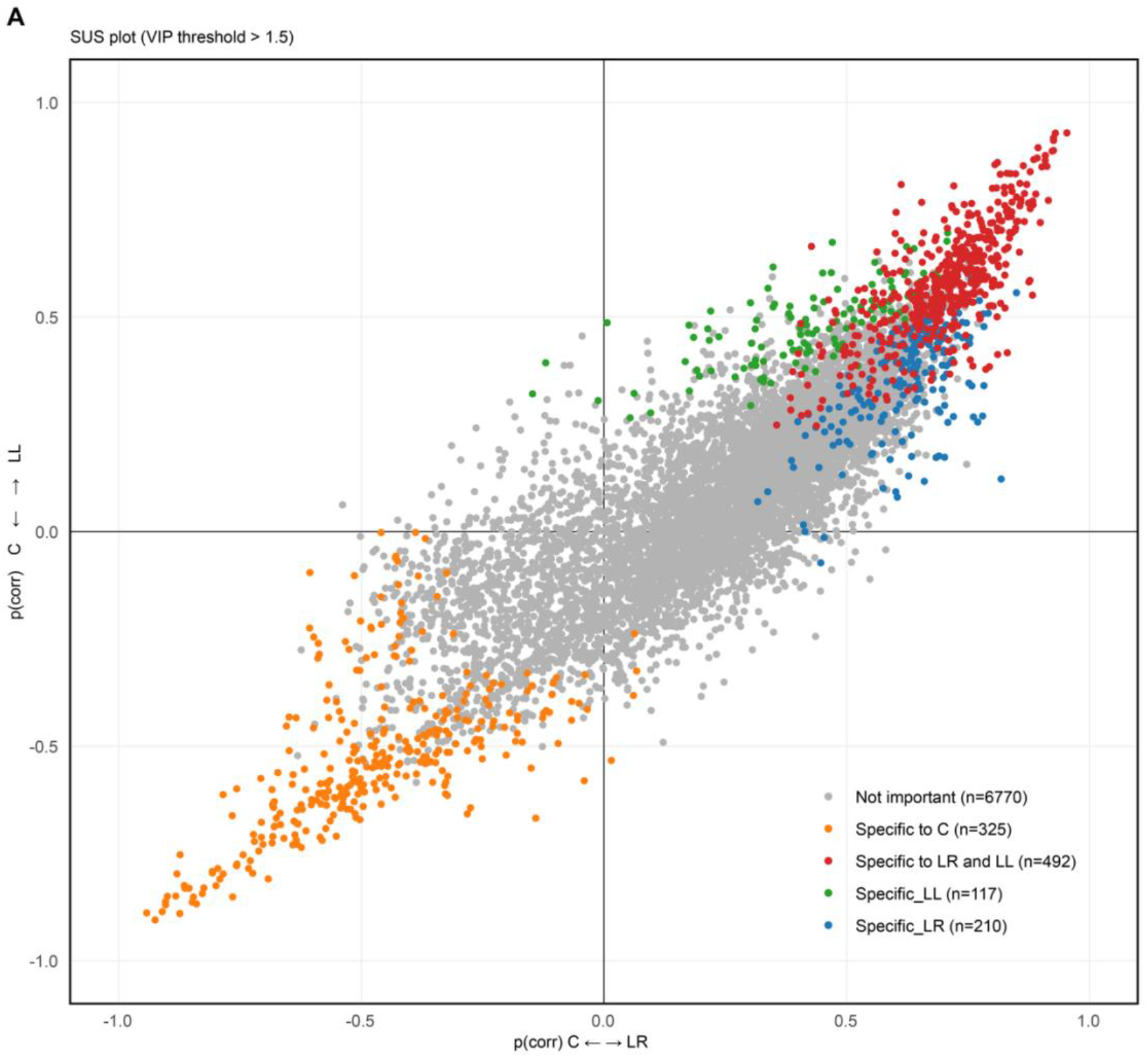
SUS plot showing shared and region-specific protein signatures. The SUS plot compares the proteins contributing to the model separating LR vs. C (x-axis; OPLS-DA, 1 predictive + 2 orthogonal components; R^2^Y=0.993, Q^2^=0.89, pQ^2^<0.01) with that of the model separating LL vs. C (y-axis; OPLS-DA, 1 predictive + 2 orthogonal components; R^2^Y=0.993, Q^2^=0.88, pQ^2^<0.01). Each point represents one protein group plotted according to its p(corr) values for the LR/C and LL/C comparisons. Protein groups passing the Variable Importance in Projection (VIP) > 1.5 threshold are colored according to their regional specificity: central ChP-specific (orange), shared between LR and LL ChP (red), LL ChP-specific (green), or LR ChP-specific (blue). Protein groups not passing these thresholds are shown in grey.

**Figure S6.**
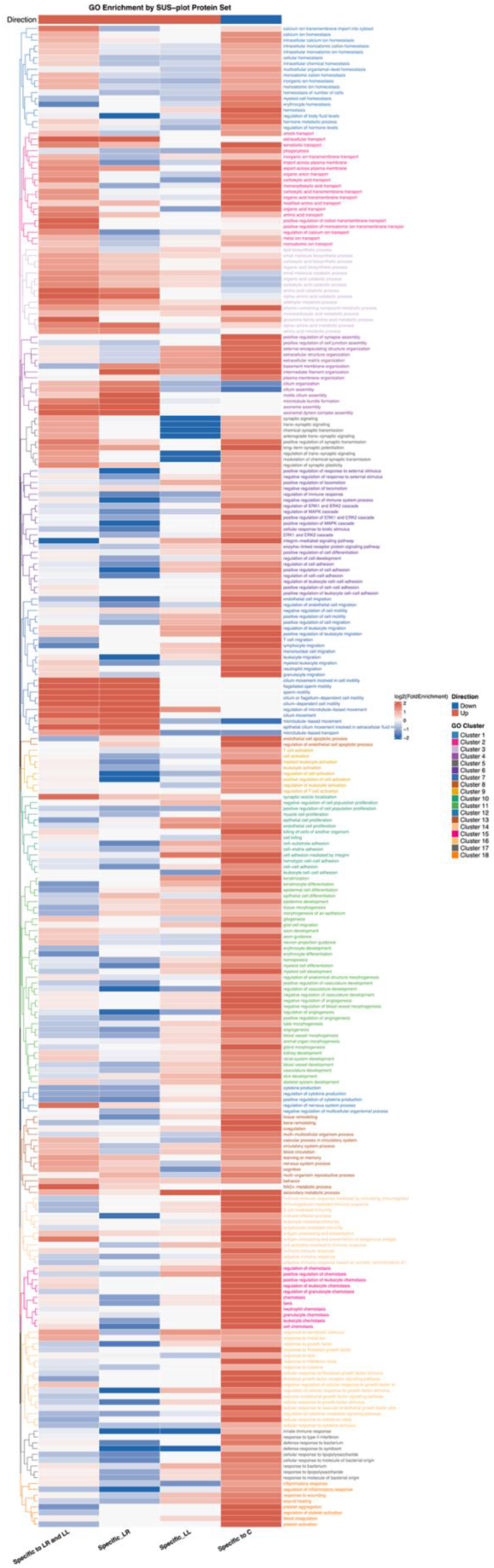
GO enrichment heatmap of SUS-defined protein sets. Heatmap displaying log2(Fold Enrichment) of significantly enriched Gene Ontology Biological Process (GO-BP) terms (Benjamini–Hochberg adjusted p < 0.01 in at least one group) across four SUS-plot-derived protein sets: proteins enriched in both LR and LL relative to C, proteins specific to the LR vs C, proteins specific to the LL vs C, and proteins enriched in C. Red indicates over-representation; white and blue cells reflect GO terms not enriched in a given protein set. Rows are clustered by semantic similarity between GO terms with dendrogram branch colors indicating cluster membership.

## Box1.

**Troubleshooting**

**Box1. Troubleshooting**
| Problem | Possible cause | Suggested solution |
| --- | --- | --- |
| **Tissue slide preparation** |  |  |
| Bubbles appear on the PEN membrane during slide preparation. | Bubble formation or membrane distortion can occur during slide preparation, especially during heat-induced or protease-induced epitope retrieval on PEN membrane slides. | Avoid harsh heating conditions when possible and carefully inspect the membrane before LMD. For FFPE samples requiring antigen retrieval, consider using a glycerol-supplemented HIER buffer, as described by Nordmann et al., to stabilize the PEN membrane and reduce distortion. Minor trapped gas or bubbles may be carefully removed and the membrane resealed. |
| Tissue section detaches during fixation and staining. | Some tissues, organoids, very soft tissues, or nervous tissue sections can be difficult to adhere to the PEN membrane. | Consider applying UV treatment, 30 min under a UV hood, or polylysine treatment prior to tissue deposition to improve tissue attachment to the PEN membrane. More information can be found in the Zeiss Protein Handling book: https://www.gu.se/sites/default/files/2020-03/Protein%20handling.pdf |
| **Microdissection** |  |  |
| Tissue pieces fly in different directions during cutting. | Insufficient attachment of the tissue to the PEN membrane. | See recommendations above for improving tissue adhesion, including UV treatment or polylysine coating prior to tissue deposition. Reduce laser power and aperture settings. |
| Cutting line does not follow the ROI drawing. | Laser misalignment between the ROI drawn on screen and the actual laser cutting path. | Perform laser calibration using Laser > Calibrate. The system cuts four calibration crosses; click the center of each cross and validate the calibration. Confirm alignment with a test cut, for example an S-shaped line on an empty membrane area. The laser calibration need to be performed for each objectives separately. |
| Tissue is very hard to cut. Laser has little to no effect, or only burns the PEN membrane rather than cutting the tissue. | Possible residual contamination from OCT embedding medium or paraffin. | Ensure that all traces of embedding material have been removed during slide preparation. OCT is water-soluble; include a water washing step after fixation or staining to remove OCT smears. For FFPE samples, deparaffinization with heating and xylene washes is mandatory before staining. |
| Burn marks: the cut becomes brown or dark around the cutting line. | Laser intensity is too high, causing local burning of the tissue or PEN membrane. | Reconsider the laser settings. Start with lower settings, for example Power 25, Aperture 1-2, and Speed 10, then re-optimize the cut quality. |
| Tissue does not fall after cutting. | The ROI may not be completely cut. | Adjust cutting Power, Speed, and Aperture. Use "Middle Pulse" mode and consider increasing the number of middle points to help detach the tissue. |
| Tissue remains adhesive to the slide around the ROI. | The tissue can remain stuck on the slide or around the ROI, potentially due to surface adhesion and/or electrostatic interactions. | Check the slide after cutting, as the tissue may remain attached around the ROI. If needed, manually detach the ROI using the Move + Cut mode to pulse the tissue piece. Perform fixation/staining and ensure complete drying to reduce surface interactions and improve tissue release. Consider adding a bar-type ionizer on the stage plate if the problem persists. |
| Cut tissue disappears or never reaches the tube. | Tissue travel may be affected by electrostatic interactions or misalignment of the stage/tube holder. | Recalibrate the stage and tube holder using the reference hole to ensure correct centering. Small pieces can be strongly affected by electrostatic interactions during travel. Consider using 0.5 mL tubes to increase the receiving area. Add a small drop of water, approximately 10-20 µL, in the cap to help capture and direct the tissue. After collection, add acetonitrile to transfer the tissue from the cap to the bottom of the tube, then dry the sample in the SpeedVac before proceeding with sample preparation. |
| **Sample preparation** |  |  |
| Loss of sample prior to sample preparation. | Tissue may stick to the side of the tube or remain in the cap. | Use 100 µL of 100% acetonitrile to transfer the tissue from the cap to the bottom of the tube. After SpeedVac drying, check recovery with a stereo microscope to ensure that the tissue has reached the bottom of the tube before proceeding with sample preparation. |
| Liquid does not pass through the SDB-RPS plug during desalting. | One air bubble can trap between the SDB-RPS disk and the liquid, preventing liquid from passing through the membrane, especially during the aqueous phase. | Centrifuge slightly faster during the first spin, around 100 × g, to prime the liquid through the plug, then immediately reduce the speed for the rest of the centrifugation run. Alternatively, briefly shake or tap the tube by hand to bring the liquid down onto the plug and help initiate flow before centrifugation. |
| Eluted sample after SDB-RPS desalting is difficult to dry in the SpeedVac. | Residual detergent or chemical contamination in the eluate. | Ensure that the SDB-RPS plug has been properly washed with both aqueous and organic wash solutions to efficiently remove DDM and other contaminants before elution. |
| **Mass spectrometry** |  |  |
| Peptide retention times are delayed or compressed near the end of the LC gradient. | Strong DDM or other chemical contamination affecting LC chromatography. | Ensure proper SDB-RPS washing with both aqueous and organic phases to remove DDM. Run LC blanks or washing gradients between runs to prevent DDM accumulation on the precolumn. Strong DDM contamination can impair peptide binding to the precolumn and affect chromatographic performance. Check for DDM-related signals in the TIC using the expected DDM molecular mass as a reference (e.g. m/z 528.34 and m/z 1021.62), as illustrated in Figure S1. |
| Low peptide recovery or low number of protein identifications, with poor TIC/MS1 signal. | Insufficient collected material, incomplete lysis or digestion, sample loss during transfer, or inefficient peptide recovery from the SDB-RPS plug. | Confirm the number or area of collected cells, for example targeting 50,000-100,000 µm<sup>2</sup> for a pilot experiment. Check tissue recovery under a stereomicroscope, ensure complete tissue lysis and digestion, and monitor digestion efficiency by checking the percentage of missed cleavages. Verify the presence of digestive-enzyme autolysis peptides at the expected retention time and abundance as an additional control for digestion performance. Verify that the SDB-RPS plug is perfectly flat, well sealed, properly conditioned, and efficiently eluted; verify that the elution buffer is close to pH 10. |
| Low peptide recovery or low number of protein identifications, with good TIC/MS1 signal. | Chemical contamination, poor peptide quality, or inefficient or absent decrosslinking in FFPE samples. | Ensure correct decrosslinking for FFPE samples, avoid excessive fixation with paraformaldehyde, and avoid non-reversible fixation methods such as glutaraldehyde. Check whether the TIC signal is dominated by non-peptide contaminants rather than true peptide signal. Check for potential polymer patterns in the TIC/MS1 chromatogram. |

**Supplementary Table 1.**

| MS1 | MS2 | Precursor<br>Window range<br>(m/z) | Number | Average<br>Window<br>Size (m/z) | Cycle time<br>(S) |
| --- | --- | --- | --- | --- | --- |
| 60 K | 30k | 430–626 | 56 | 3.5 | 3.712 |
| 60 K | 60k | 430–626 | 28 | 7 | 3.712 |
| 60 K | 120k | 430–626 | 14 | 14 | 3.712 |
| 60 K | 240k | 430–626 | 7 | 28 | 3.712 |
| 60 K | 30k | 418–670 | 56 | 4.5 | 3.712 |
| 60 K | 60k | 418–670 | 28 | 9 | 3.712 |
| 60 K | 120k | 418–670 | 14 | 18 | 3.712 |
| 60 K | 240k | 418–670 | 7 | 36 | 3.712 |
| 60 K | 30k | 416–808 | 56 | 5.5 | 3.712 |
| 60 K | 60k | 416–808 | 28 | 11 | 3.712 |
| 60 K | 120k | 416–808 | 14 | 22 | 3.712 |
| 60 K | 240k | 416–808 | 7 | 44 | 3.712 |
| 60 K | 30k | 416–976 | 56 | 10 | 3.712 |
| 60 K | 60k | 416–976 | 28 | 20 | 3.712 |
| 60 K | 120k | 416–976 | 14 | 40 | 3.712 |
| 60 K | 240k | 416–976 | 7 | 80 | 3.712 |

